# Understanding responses to multi-electrode epiretinal stimulation using a biophysical model

**DOI:** 10.1101/2024.08.20.608829

**Authors:** Ramandeep S. Vilkhu, Praful K. Vasireddy, Kathleen E. Kish, Alex R. Gogliettino, Amrith Lotlikar, Pawel Hottowy, Wladyslaw Dabrowski, Alexander Sher, Alan M. Litke, Subhasish Mitra, E.J. Chichilnisky

## Abstract

**Objective:** Neural interfaces are designed to evoke specific patterns of electrical activity in populations of neurons by stimulating with many electrodes. However, currents passed simultaneously through multiple electrodes often combine nonlinearly to drive neural responses, making evoked responses difficult to predict and control. This response nonlinearity could arise from the interaction of many excitable sites in each cell, any of which can produce a spike. However, this multi-site activation hypothesis is difficult to verify experimentally.

**Approach:** We developed a biophysical model to study retinal ganglion cell (RGC) responses to multi-electrode stimulation and validated it using data collected from *ex vivo* preparations of the macaque retina using a microelectrode array (512 electrodes; 30µm pitch; 10µm diameter).

**Results:** First, the model was validated by using it to reproduce essential empirical findings from single-electrode recording and stimulation, including recorded spike voltage waveforms at multiple locations and sigmoidal responses to injected current. Then, stimulation with two electrodes was modeled to test how the positioning of the electrodes relative to the cell affected the degree of response nonlinearity. Currents passed through pairs of electrodes positioned near the cell body or far from the axon (>40 µm) exhibited approximately linear summation in evoking spikes. Currents passed through pairs of electrodes close to the axon summed linearly when their locations along the axon were similar, and nonlinearly otherwise. Over a range of electrode placements, several distinct, localized spike initiation sites were observed, and the number of these sites covaried with the degree of response nonlinearity. Similar trends were observed for three-electrode stimuli. All of these trends in the simulation were consistent with experimental observations. *Significance*. These findings support the multi-site activation hypothesis for nonlinear activation of neurons, providing a biophysical interpretation of previous experimental results and potentially enabling more efficient use of multi-electrode stimuli in future neural implants.

## Introduction

Electronic neural implants can enable novel treatment strategies for intractable nervous system disorders. An important example is epiretinal implants for treating photoreceptor degenerative diseases such as *retinitis pigmentosa* and age-related macular degeneration [1–3]. Epiretinal implants electrically stimulate on the surface of the retina to evoke patterns of activity in retinal ganglion cells (RGCs), thereby sending artificial visual signals to the brain. Although modern epiretinal implants have been able to produce coarse light perception, they have not been able to create formed visual percepts that correspond well to the visual environment. One reason for this difficulty is that currents passed through multiple electrodes simultaneously often sum nonlinearly to drive RGC responses, making the evoked responses difficult to predict and control [4]. Therefore, to enable the more effective use of multi-electrode stimulation it is likely necessary to quantitatively understand the biophysical basis of this nonlinearity.

One hypothesis to explain the nonlinear responses of RGCs is that stimulation currents from many electrodes interact with many excitable sites along a cell, any of which can produce a spike [4]. Thus, while currents likely combine linearly to drive activation at any particular site, the overall neural response is a nonlinear “OR” combination across all the sites [4–6]. However, this *multi-site activation hypothesis* has not been biophysically tested and previous studies reported varying degrees of nonlinear summation, the basis of which is still not well understood. Notably, a biophysical abstraction, the activating function, has previously been used to predict RGC activation [7]. However, its utility in predicting the precise location of spike initiation, which is needed to probe multi-site activation, is limited for realistic RGC geometries [8,9].

Here, we developed a biophysical model to test whether the multi-site activation hypothesis can explain nonlinear responses to multi-electrode stimulation [9,10]. The model was validated against experimentally recorded responses of individual RGCs to electrical stimulation, using data collected from *ex vivo* preparations of the macaque retina [11,12]. We observed that varying degrees of response nonlinearity could be explained by variability in the geometric positioning of the electrodes relative to the cell. Then, by using the model to probe the locus of spike initiation, distinct sites of activation were revealed. Over a range of electrode placements, the number of observed activation sites covaried with the degree of response nonlinearity. This provided biophysical support for the multi-site activation hypothesis and a quantitative understanding of geometric factors causing varying degrees of nonlinear current summation. These insights could allow for better prediction of the expected response linearity, enabling the more effective use of multi-electrode stimulation in future neural implants.

## Methods

### Biophysical model

The biophysical model of a spiking RGC was developed in the NEURON [10] simulation environment, with details given below. The model is described schematically in **Fig. 1**. All code used to build the RGC model is online and available at: https://github.com/ramanvilkhu/rgc_simulation_multielectrode

**Figure 1.**
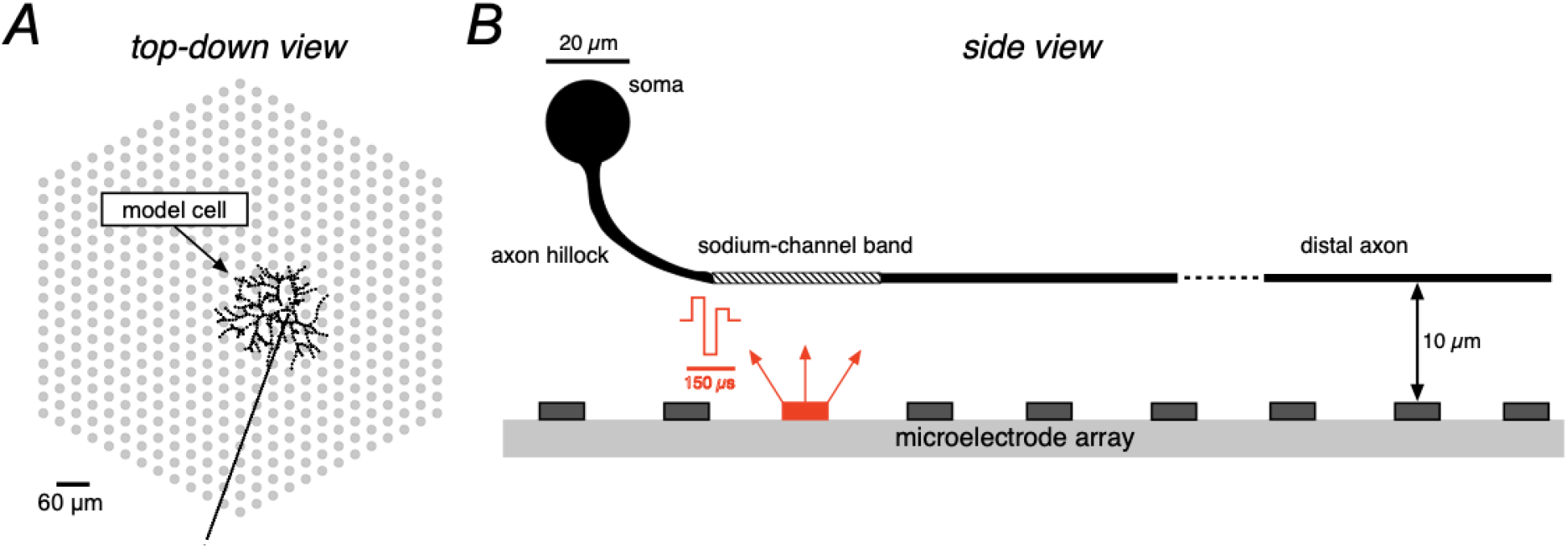
Visualization of the simulation setup. ***A***, Single modeled parasol cell, traced in black, with realistic dendritic structure taken from [13] and a thin linear axon against the simulated microelectrode array (MEA) with electrode diameter and pitch of 10µm and 30µm, respectively. ***B***, Side-view of the model cell showing the various modeled compartments and the bend in the RGC axon as it extends from the soma to the nerve fiber layer, where the planar MEA is positioned, matching experimental conditions (see Methods). Charge-balanced triphasic stimulus with relative phase amplitudes of (2:-3:1) supplied through a single electrode (red). Dendrites not shown.

### Biophysics and morphology

The biophysical properties of the RGC cable model used were adapted from previous work [9] with two modifications. First, the temperature of the simulation environment was set to 35°C, matching that of the experimental setup, and the membrane dynamics were adjusted according to the Q10 factors detailed in previous work [14]. Second, the narrow axon compartment was removed and the remaining compartment diameters were increased, which allowed for more stable model behavior when simulating electrical stimulation with multiple electrodes. The remaining parameters were not changed, and all properties are described below.

The cytoplasmic (axial) resistivity was 143.2 Ω cm and the membrane capacitance was 1 µF cm^-2^ across all compartments. The differential equation governing membrane voltage is given in Equation 1.

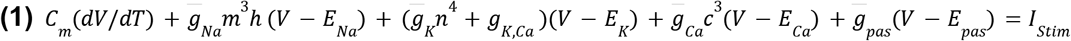

The channel state variables (*m, h, n, c*) for the voltage-gated ion channels (Na^+^, K^+^, Ca^2+^) are governed by equations of the form shown in Equation 2. The voltage-dependent rate equations for each of these state variables are shown in Table 1, taken from previous work (see [14], table 2).

**Table 1:** Rate constants for voltage-gated ion channels.

| Sodium (Na<sup>+</sup>) | Potassium (K<sup>+</sup>) | Calcium (Ca<sup>2+</sup>) |
| --- | --- | --- |
| $\alpha_m = \frac{-2.725(V+35)}{e^{-0.1(V+35)} - 1}$ | $\alpha_n = \frac{-0.09575(V+37)}{e^{-0.1(V+37)} - 1}$ | $\alpha_c = \frac{-1.362(V+13)}{e^{-0.1(V+13)} - 1}$ |
| $\beta_m = 90.83e^{\frac{-(V+60)}{20}}$ | $\beta_n = 1.915e^{\frac{-(V+47)}{80}}$ | $\beta_c = 45.41e^{\frac{-(V+38)}{18}}$ |
| $\alpha_h = 1.817e^{\frac{-(V+52)}{20}}$ | | |
| $\beta_h = \frac{27.25}{1+e^{-0.1(V+22)}}$ | | |

**Table 2:** Nerst potential for ion channels (mV).

| E<sub>Na</sub> | E<sub>K</sub> | E<sub>Ca</sub> | E<sub>Pas</sub> |
| --- | --- | --- | --- |
| 60.60 | -101.34 | $\frac{RT}{2F} \ln\left(\frac{[Ca^{2+}]_{outside}}{[Ca^{2+}]_{inside}}\right)$ | -64.58 |

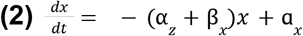

Unlike the other ion channels, the calcium-activated potassium channel (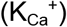) is ligand-gated according to Equation 3. The calcium ion concentration is driven by a pump mechanism, given by Equation 4, where *F* is Faraday’s constant (96,485 °C) and *r* is the depth (0.1 µm) at which the calcium ion concentration is measured.

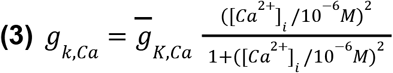

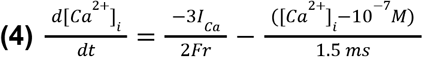

The Nernst potential for each ion is defined in Table 2. These potentials are constant during the spike for all ions, except calcium.

The RGC is divided into a total of five distinct regions, three of which are distinct sections of the axon. Table 3 outlines the maximum ion channel conductance values for each region and Table 4 shows the physical characteristics of each region (e.g. length, diameter). These values were adapted from [9]: the diameters of the compartments were modified slightly to enable more robust model behavior in response to multi-electrode stimulation. Notably, the maximum ion channel conductance values of each region were within the range of previously reported values [8,15].

**Table 3:** Maximum ion channel conductance values at each compartment (mS cm^-2^).

| | $\bar{g}_{Na}$ | $\bar{g}_K$ | $\bar{g}_{Ca}$ | $\bar{g}_{K,Ca}$ | $\bar{g}_{Pas}$ |
| --- | --- | --- | --- | --- | --- |
| dendrites | 60 | 35 | 1 | 0.17 | 0.1 |
| soma | 60 | 35 | 0.75 | 0.17 | 0.1 |
| axon hillock | 150 | 90 | 0.75 | 0.17 | 0.1 |
| sodium channel band | 420 | 250 | 0.75 | 0.11 | 0.1 |
| distal axon | 100 | 50 | 0.75 | 0.2 | 0.1 |

**Table 4:** Morphological details of compartments (length and diameter) (µm).

|  | length | diameter |
| --- | --- | --- |
| dendrites | from NeuronDB [13] |  |
| soma | 20 | 20 |
| axon hillock | 40 | 4 |
| sodium channel band | 40 | $4 \rightarrow 1.5$ |
| distal axon | 2970 | 1.5 |

The dynamics were encoded into a multi-compartment cable model, which was implemented in Python using NEURON. The model was discretized into 5 µm segments and the time resolution of the solver was set to 0.005 ms time steps, matching the values shown in previous work to maximize computational efficiency [9].

### Extracellular electrical stimulation

The extracellular tissue medium was approximated as an isotropic ohmic conductor. A disk electrode acting as a current source with ground located at infinity was placed below (∼10 µm) the modeled RGC. Since the extracellular domain was isotropic, the extracellular voltage at each point in space was calculated in a similar fashion to previous work [16]. Shown in Equation 5:

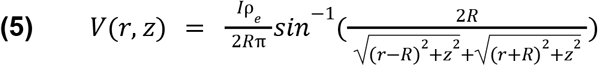

where *r* and *z* are the radial and axial displacement from the center of the disk for *z* ≠ 0. *R* is the radius of the disk electrode (5 µm), *I* is the stimulation current, and ρ_e_ is the extracellular resistivity (1000Ω *cm*) [8]. Notably, a large range of resistivity values for the ganglion cell layer is reported in the literature (between 500 and 7900 Ω *cm*) [17–19]. In this work, 1000 Ω *cm*, which is used in several other studies, yielded the best match to the experimental data [8]. The electrode array was placed 10 µm below the nerve-fiber layer (shown in **Fig. 1**), which was chosen to match the recorded voltage amplitudes seen in the data and was similar to previous estimates [20].

Electrical stimuli consisted of triphasic charge-balanced pulses with relative current amplitudes 2:-3:1 and duration of 50 μs per phase (150 μs total), matching experimental stimulation parameters. The *activation threshold* was defined as the lowest current amplitude required to elicit a spike. Similarly, for multi-electrode stimulation, for each fixed ratio of current levels passed through the electrodes, the smallest current amplitude combination that produced a spike was identified as the *activation threshold*.

### Stochastic spiking

To model stochastic spiking [21], a Gaussian noise membrane current was added to the total ionic membrane currents as described in previous work [22]. The noise current was on the order of microamps and was proportional to the square root of the number of sodium channels. It was independently added to each segment *n* in the model according to Equation 6:

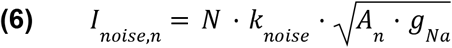

where *N* is a Gaussian noise draw (µ = 0; σ = 1) at each time step of the NEURON solver (0.005 ms), *k*_*noise*_ was a constant set to 0.0004 in order to best match experimental data, *A*_*n*_ denotes the membrane area in *cm*^2^, and *g*_*Na*_ is the maximum sodium conductance at that segment.

### Determination of threshold and tracking the locus of spike initiation

To determine if the model cell spiked, the membrane potential at the far distal end of the model axon was monitored. If the membrane potential crossed 0 mV within 10 ms of the stimulus onset, the model cell was considered to have spiked. For each fixed ratio of current levels passed through the multiple electrodes, the smallest current amplitude combination that produced a spike was taken as the *activation threshold*.

The segment in which the membrane potential first crossed 0 mV after completion of the stimulus pulse was taken as the site of spike initiation. Occasionally, during multi-electrode stimulation, a continuous band of 2-3 neighboring segments all had the same timing of membrane zero-crossing within the time resolution of the simulation. In these cases, the exact location was indistinguishable between these segments, so the locus of spike initiation was taken to be the center of this region. Information about the locus of activation was further encoded in the color of the data in Fig. 3, Fig. 4, and Fig. 7. A shift in the color illustrated a shift in the locus of spike initiation.

**Figure 2.**
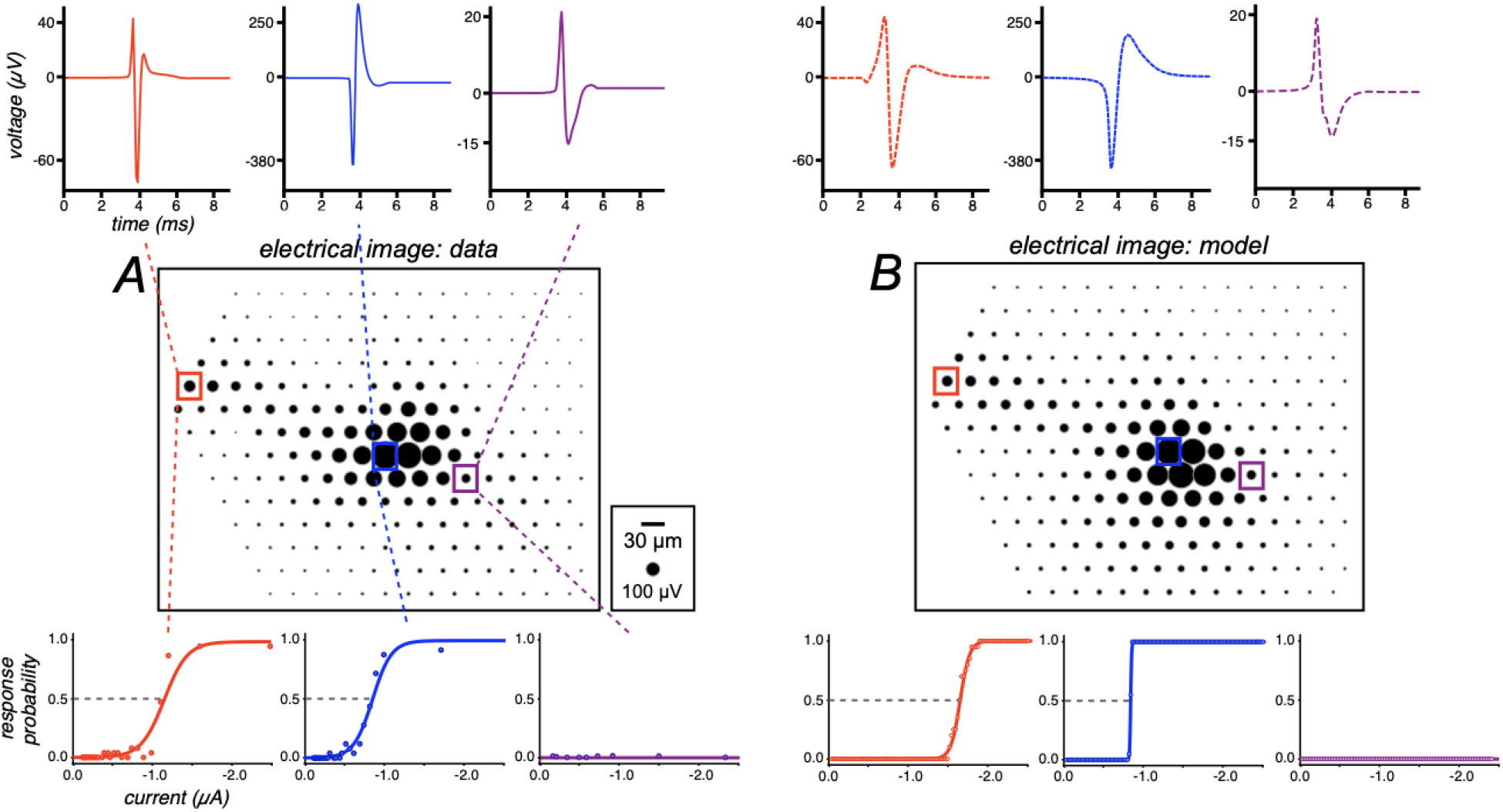
Single-electrode recording and stimulation of a single parasol RGC with a large-scale electrode array. ***A***, Experimental electrical image (EI) from a parasol cell. Black circle area is proportional to recorded signal strength. Top, Examples of recorded axonal (red), somatic (blue), and dendritic (purple) spike waveforms, detected at electrodes marked by the respective colored square. Bottom, Electrical activation as a function of current level at the selected electrodes from the distinct cellular compartments. **B**, Similar to **A**, but for a modeled parasol cell. The spatial correlation coefficient between the simulated and empirical EI for this cell was 0.73 (see Methods).

**Figure 3.**
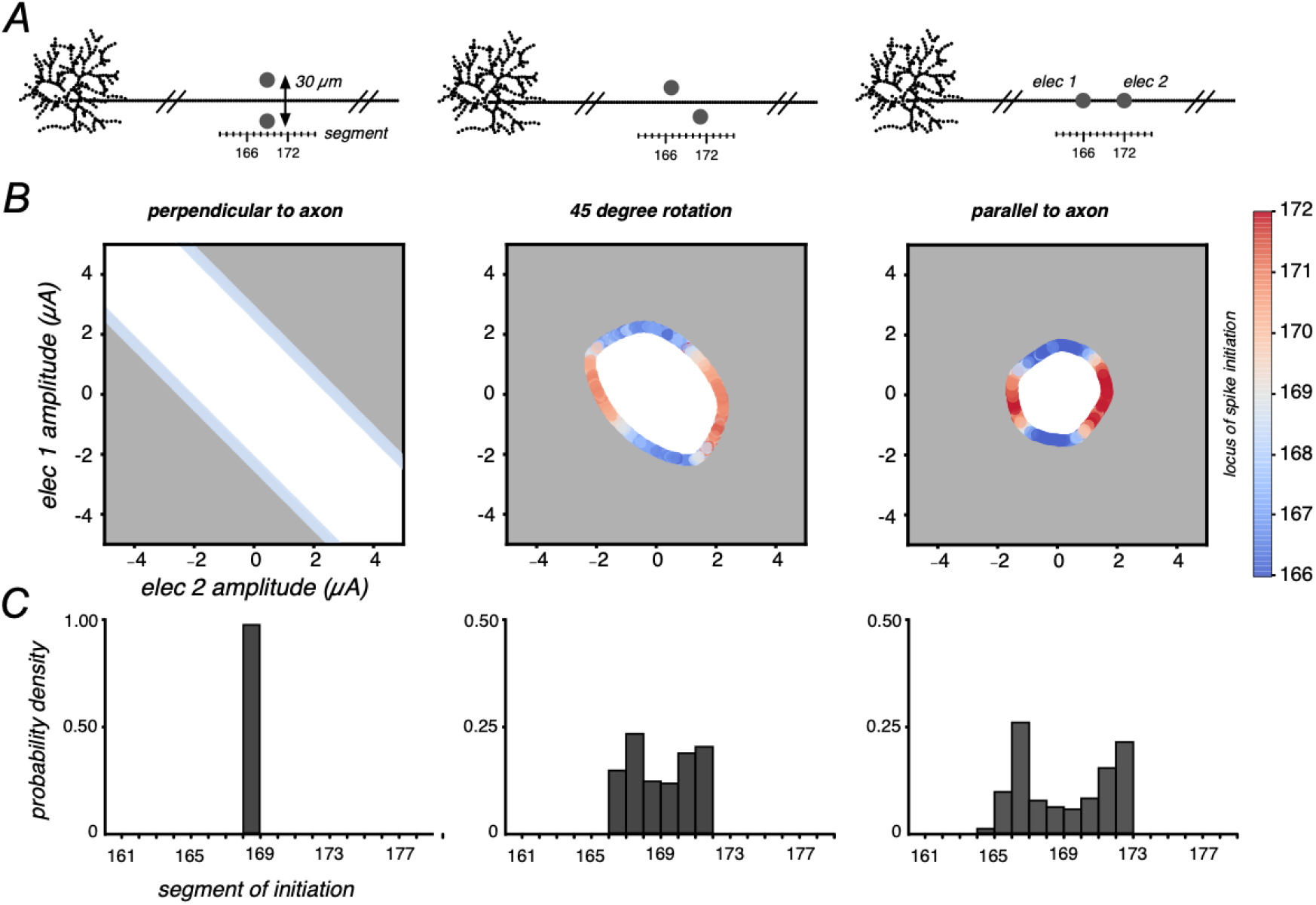
Responses to two electrode stimulation, orientation effects. ***A***, Visualization of the two electrodes, separated by 30 µm, positioned at various orientations relative to the distal axon of the model cell. The model cell is discretized into 5 µm length segments (see Methods). ***B***, Collection of activation thresholds in the two-dimensional space of current levels, colored by segment of spike initiation. Grey region denotes current combinations above threshold that cause spiking. ***C***, Distribution of spike initiation locations for the collection of current combinations causing activation that were colored in the middle panel (see Methods).

**Figure 4.**
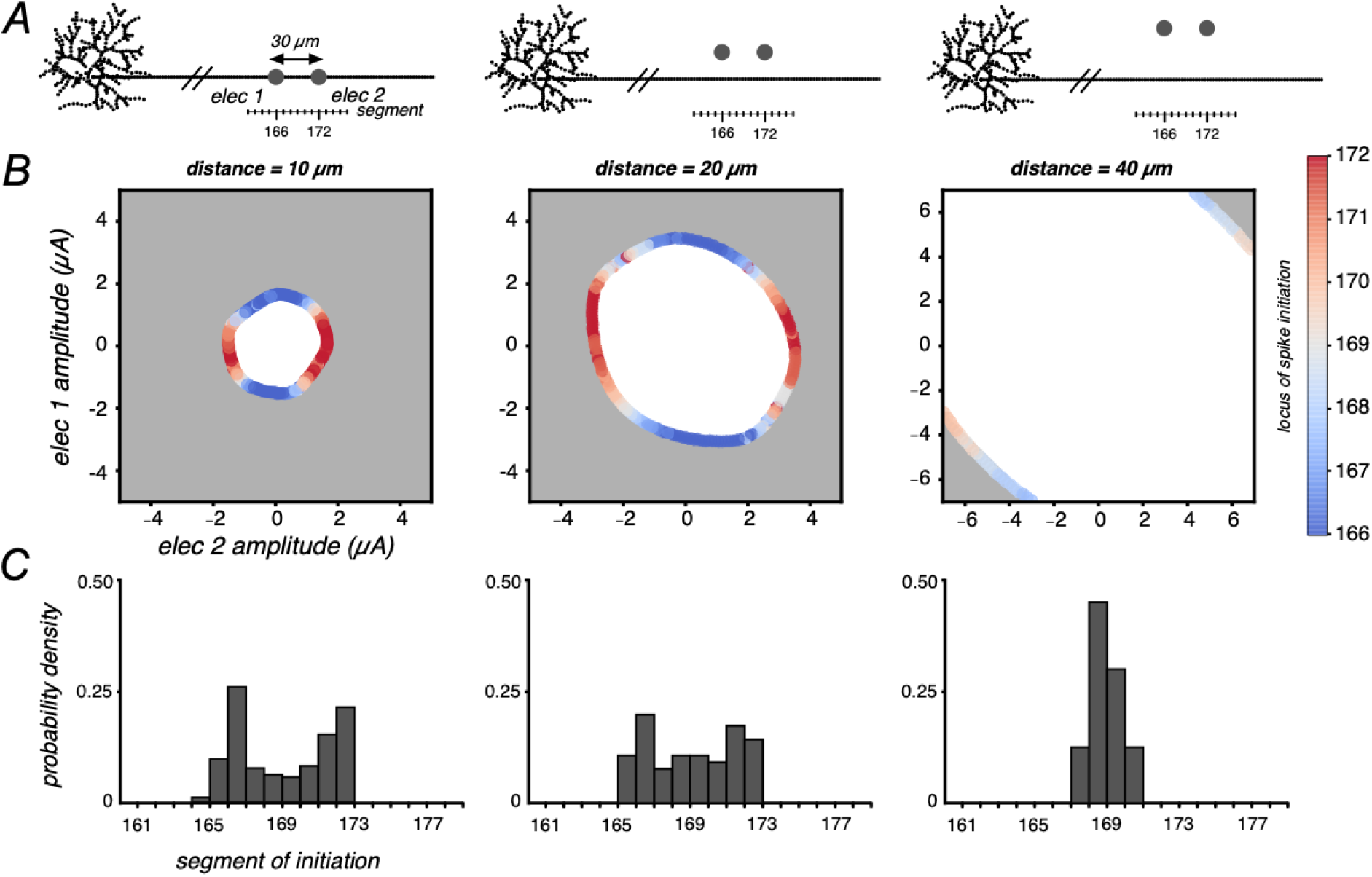
Responses to two electrode stimulation, distance effects. ***A***, Visualization of the two electrodes, separated by 30 µm, oriented parallel to the distal axon, and positioned at various distances from the model cell. The model cell is discretized into 5 µm length segments (see Methods). ***B***, Collection of activation thresholds in the two-dimensional space of current levels, colored by segment of spike initiation. Grey region denotes current combinations above threshold that cause spiking. ***C***, Distribution of spike initiation locations for the collection of current combinations causing activation that were colored in the middle panel (see Methods).

### Nonlinearity index

A nonlinearity index was calculated to assess the degree of nonlinear current summation present in modeled responses to stimulation using two electrodes. For various placements of a pair of electrodes across the somatic and dendritic regions of the cell, activation thresholds were computed for various fixed ratios of current passed through the two electrodes. An ellipse was fitted to the collection of activation thresholds in the two-dimensional space of current levels on the two electrodes, using a least-squares minimization approach. The nonlinearity index was defined as the ratio of the lengths of the minor and major axes of the fitted ellipse. The resulting value ranged between 0 (corresponding to an infinitely stretched ellipse fitting linear data) and 1 (corresponding to a circle fitting highly nonlinear data). This procedure was repeated for four orientations of the electrode pair (parallel, perpendicular, and two 45 degree rotations) relative to the cell, and the maximum value across rotations was used as a summary.

### Experimental setup

A custom 512-electrode system [12] was used to stimulate and record from RGCs in isolated rhesus macaque monkey (*Macaca mulatta*) retinas. Eyes were obtained from terminally anesthetized animals euthanized during the course of research performed by other laboratories. All procedures were performed in accordance with institutional and national guidelines and regulations. Briefly, the eyes were hemisected in room light following enucleation, the vitreous was removed and the posterior portion of the eye was kept in darkness in warm (35°C), oxygenated, bicarbonate buffered Ames solution (Sigma). Then, patches of retina ∼3 mm on a side were isolated under infrared light, placed RGC side down on the multielectrode array, and superfused with Ames solution. The microelectrode array was composed of 512 electrodes (diameter, 10 µm) with 30 µm pitch, covering an area of 0.43 mm^2^. Within an experiment, platinization of the electrodes produced relatively uniform noise, approximately 6% (standard deviation) across electrodes. A platinum wire encircling the recording chamber (∼1 cm diameter) served as the distant return electrode. Voltage recordings were band-pass filtered between 43 and 5,000 Hz and sampled at 20 kHz. Spikes from individual RGCs in the voltage recordings were identified and sorted using standard techniques [23].

### Visual stimulation and cell type classification

To identify the retinal ganglion cell types recorded, the retina was visually stimulated with a dynamic white noise stimulus, and the spike-triggered average (STA) stimulus was computed for each RGC, as previously described [24,25]. The STA summarizes the spatial, temporal, and chromatic properties of light response. In particular, clustering on the spatial (receptive field size) and temporal (time course) components of the STA was performed to identify distinct cell types, as previously described [26]. Analysis focused on ON and OFF parasol RGCs due to the high SNR of their recorded spikes, which was useful for reliable spike sorting in the presence of electrical artifacts (see below).

### Electrical image

The electrical image (EI) represents the average spatiotemporal pattern of voltage deflections produced on each electrode of the array during a spike from a given cell [11]. Electrical images were calculated from data recorded during visual stimulation and served as spatiotemporal templates for the spike waveforms of the cells to be detected during electrical stimulation.

The spatial positions of relevant cell compartments (axon and soma) of a recorded cell relative to the electrode array were estimated using the electrical image. For each recorded cell, the shape of the spike waveform from the EI on an electrode was used for compartment identification. A triphasic waveform was taken to indicate recording from the cell axon, a biphasic waveform with a positive first phase was taken to indicate recording from the cell dendrite, and a biphasic waveform with a negative first phase was taken to indicate recording from the cell soma [11].

A spatial correlation metric was used to quantitatively compare modeled and empirical EIs without over-penalizing known discrepancies in action potential dynamics (see Discussion). First, the total power of the voltage associated with a spike (sum of squared voltage values over time) was calculated on each electrode for both the modeled and empirical data. Then, the Pearson correlation coefficient was computed for this value across electrodes.

### Electrical stimulation

Electrical stimulation was provided through one or more electrodes while recording RGC activity from all electrodes. Three types of stimulation patterns were tested: single-electrode stimulation, two-electrode stimulation, and three-electrode stimulation. Single-electrode stimulation consisted of a charge-balanced, triphasic pulse passed through one electrode. The negative polarity stimulus consisted of a triphasic pulse with anodal/cathodal/anodal phases with relative current amplitudes 2:-3:1 and duration of 50 μs per phase (150 μs total). The positive polarity stimulus was similar but with the polarity of the phases flipped to be cathodal/anodal/cathodal. Single-electrode stimulation was delivered in 25 repeated trials at 40 logarithmically spaced current amplitudes (10% increments) between 0.1 and 4 μA. Two-electrode and three-electrode stimulation consisted of triphasic, charge-balanced current simultaneously passed through two or three adjacent electrodes, respectively. The stimulating electrodes were chosen based on the highest SNR of recorded spikes from the target cell as well as the geometric positioning of the electrodes relative to the target cell. Stimulation was supplied for 20 trials at 20 linearly spaced current amplitudes between -1.8 and 1.8 μA. As a result, for two-electrode and three-electrode stimulation, 400 and 8000 unique current combinations were tested, respectively. The ordering of the stimulation patterns was chosen pseudo-randomly, restricted so that each successive stimulating electrode or group of electrodes was far from the previous and subsequent stimulating electrodes, to avoid stimulating the same cell(s) in rapid succession.

### Responses to electrical stimulation

The spikes recorded during electrical stimulation were analyzed using a custom template matching approach [27]. First, the electrical image of each cell was calculated from visual stimulation data, and served as a template for the spike waveform of each cell to be detected during electrical stimulation. In the electrical stimulation data, an automated algorithm separated spikes from the electrical artifact by grouping traces according to the artifact waveform estimate and the spike waveform of each cell analyzed. The resulting electrically elicited spike waveforms were visually inspected for sorting errors and manually corrected as needed.

For single-electrode stimulation, spike probabilities for each cell were calculated across the trials at each stimulus current amplitude, and spiking probability was modeled as a function of current amplitude by a sigmoidal relationship (Equation 1).

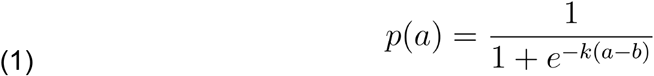

where a is the current amplitude, p(a) is the spike probability, and k and b are free parameters (representing sigmoidal slope and threshold, respectively). The fitted sigmoidal function was used to compute the experimentally observed *activation threshold*, defined as the current amplitude *a* producing 0.5 spiking probability.

For two-electrode stimulation, spike probabilities for each cell were calculated across trials for all current combinations passed through the two electrodes. Then, in the resulting 2D space of currents, 30 direction vectors starting at the origin and extending radially were computed, representing fixed current ratios through the two electrodes. For each unique current ratio, the data near the direction vector were gathered and a sigmoid was fitted to the data. The point along that vector that caused 0.5 spiking probability was denoted as the two-electrode *activation threshold*. The arrangement of the collection of activation thresholds in the two-dimensional space of current levels was examined to test linearity of summation. If currents combine linearly to drive response, then the thresholds should lie on a line (or, for pulses with positive and negative phases such as the ones used here, two lines on opposite sides of the origin).

For three-electrode stimulation, spike probabilities for each cell were calculated across trials for all current combinations passed through the three electrodes. For plotting, only combinations causing spike probability 0<p<1 were used to examine the shape of the response in the three-dimensional space of currents. Activation thresholds were not computed for three-electrode stimulation.

## Results

To investigate the mechanisms underlying RGC responses to multi-electrode stimulation, we developed a biophysical model and validated it against data from *ex vivo* preparations of the macaque retina collected with a large-scale, high-density microelectrode array (512 electrodes, 30 µm pitch, 10 µm diameter). Below, we demonstrate the model’s ability to reproduce a wide range of empirical findings, such as stereotypical voltage waveforms of recorded spikes and sigmoidal response probabilities as a function of stimulation current level. We then show that current passed through multiple electrodes simultaneously sometimes sums linearly to drive spiking activity, and sometimes sums nonlinearly, depending on the geometrical arrangement of the electrodes relative to the cell, and validate this finding with empirical data. Finally we use the model to test the hypothesis that shifts in the locus of spike initiation can explain nonlinear current summation [4].

### Modeling single-electrode stimulation and recording

The model (described schematically in **Fig. 1)** captured salient properties of large-scale extracellular voltage recording from RGCs. The modeled electrical image (EI) – the spatiotemporal voltage pattern recorded across the electrode array during a spike – closely matched the experimentally observed EI (**Fig. 2, middle**). Additionally, the modeled recording of the actional potential at the axon was triphasic, and at the soma and dendrites was biphasic with opposing polarity in the two compartments (**Fig. 2, top**). The spatial correlation coefficient between the simulated EI and empirical EI for the five cells examined was 0.82 ± 0.09 (mean ± SD). A shortcoming of the model is that the temporal dynamics of the modeled spikes were slower than what was observed experimentally (see Discussion).

Essential properties of spikes evoked by extracellular electrical stimulation were also captured by the model. Spikes were evoked using current levels similar to those used experimentally, with the lowest threshold for activation (∼1 µA) occurring near the axon initial segment, consistent with previous work (**Fig. 2, bottom**) [28,29]. Within the current range tested with single-electrode stimulation, dendritic activation and upper threshold phenomenon (lack of activation at high current levels due to the reversal of the sodium ion channel current at strong depolarizations) were not observed, although they were observed using larger currents (not shown).

The model reproduced the characteristic sigmoidal response probability as a function of current level (**Fig. 2, bottom**) [21,22]. The slopes of the modeled sigmoids varied inversely with the stimulation threshold (current level producing a spike with probability 0.5), consistent with experimental data [20]. As expected, the activation threshold in the stochastic model corresponded to the lowest current level that caused a spike in the non-stochastic model. Thus, for computational efficiency, in what follows only the non-stochastic implementation of the model was used, and the activation threshold was defined as the lowest current level that caused a spike.

### Two-electrode stimulation: linear and nonlinear responses

Previous work has shown that currents passed simultaneously through multiple electrodes can combine linearly or nonlinearly to drive RGC response [4]. A potential explanation is that currents combine nonlinearly if the electrodes target distinct spike initiation sites on the cell (see Discussion for details). Below, we first show that the model exhibits both linear and nonlinear current summation, as seen experimentally, and then probe spike initiation to test the multi-site activation hypothesis.

First, electrodes were placed close to the axon (<40 µm) and arranged at several orientations relative to the axon (**Fig. 3A**). In this case, currents combined linearly to drive firing when the pair was placed perpendicular to the axon (**Fig 3A, left**): the collection of activation thresholds formed two parallel lines in the space of electrical stimuli (**Fig. 3B, left**). In contrast, currents combined nonlinearly when the electrode pair was rotated to be more parallel to the axon (**Fig. 3A, middle, right**): the collection of thresholds formed a curved and closed shape in the space of electrical stimuli (**Fig. 3B, middle, right**). These observations are consistent with the idea that current from two electrodes combines linearly in producing activation only if they access the same activation site. To further test this idea, the sites of activation were identified as the earliest supra-threshold voltage location in the model output (see Methods). In the case of linear summation, the observed sites of activation accumulated across all current combinations at threshold were highly localized along the axon (**Fig. 3C, left**). In the cases of nonlinear summation, the locations were more widely spread along the axon, with two distinct peaks occurring at sites aligned with the two electrodes (**Fig. 3C, middle & right**). Examination of the site of activation revealed that the primary site of activation progressively shifted as the dominant current amplitude moved from one electrode to the other (**Fig. 3B, middle & right, colors**).

Second, the electrode pair was placed parallel to the axon but at varying distances from it (**Fig. 4A**). In this case, the response nonlinearity decreased (**Fig. 4B**) as the electrode pair was moved away from the axon. The currents combined approximately linearly for distances greater than 40 µm (**Fig. 4B, right**). This finding is consistent with the idea that at long distances, the radial dispersion of current from each of the two electrodes causes them to target largely overlapping regions of the axon, yielding linear summation, while at shorter distances, they target more distinct regions, yielding nonlinear summation. As in the preceding analysis, the sites of activation for the electrode placements were tracked in the model to test the multi-site activation hypothesis and similar trends emerged. In the case of linear summation, the observed region of activation across different threshold current combinations was tightly localized on the axon (**Fig. 4C, right**). In the cases of nonlinear summation, the region of activation was more widely spread, with distinct peaks appearing near the regions aligned with the two electrodes (**Fig. 4C, left & middle**). In the nonlinear cases, the primary site of activation systematically shifted as the dominant current amplitude moved from one electrode to the other (**Fig. 4B, left & middle, colors**).

Third, the electrode pair was placed near somatic and dendritic regions of the cell, with the two electrodes separated by 30 µm, and an index was computed that captures the maximum degree of nonlinearity across rotations of the electrodes (see Methods). The nonlinearity of summation near the soma and dendrites was substantially smaller than was observed near the axon (**Fig. 5**. This finding is consistent with the idea that spike initiation in these cases predominantly occurs at the axon initial segment, which is distant (>40 µm) from most somatic and dendritic electrode placements [8]. Although summation close to the AIS displayed significant nonlinearity for some rotations, stimulation with electrode pairs arranged perpendicular to the axon was linear (not shown), similar to the results observed with the distal axon (**Fig. 3**).

To experimentally verify the trends observed in simulation, the spiking behavior of parasol RGCs in response to stimulation from pairs of electrodes at various geometric placements relative to the cell was measured in *ex vivo* preparations of the macaque retina. The experimental trends broadly matched those seen in simulations. Specifically, for electrode pair placements close to the axon, currents demonstrated nonlinear summation when the electrodes were parallel to the axon (**Fig. 6A**) and more linear summation when they were arranged perpendicular to the axon (**Fig. 6B**). Electrode pairs placed in a parallel arrangement but far (>40 µm) from the axon demonstrated no spiking within the experimentally tested current range. Electrode pair positions over the somatic and dendritic regions of the cell demonstrated more linear current summation (**Fig. 6C**). Overall, the distinct shapes of the collection of activation thresholds also approximately matched those observed in previous experiments [4], supporting the notion that geometric variability produced the varying amounts of nonlinearity observed in that work.

**Figure 5.**
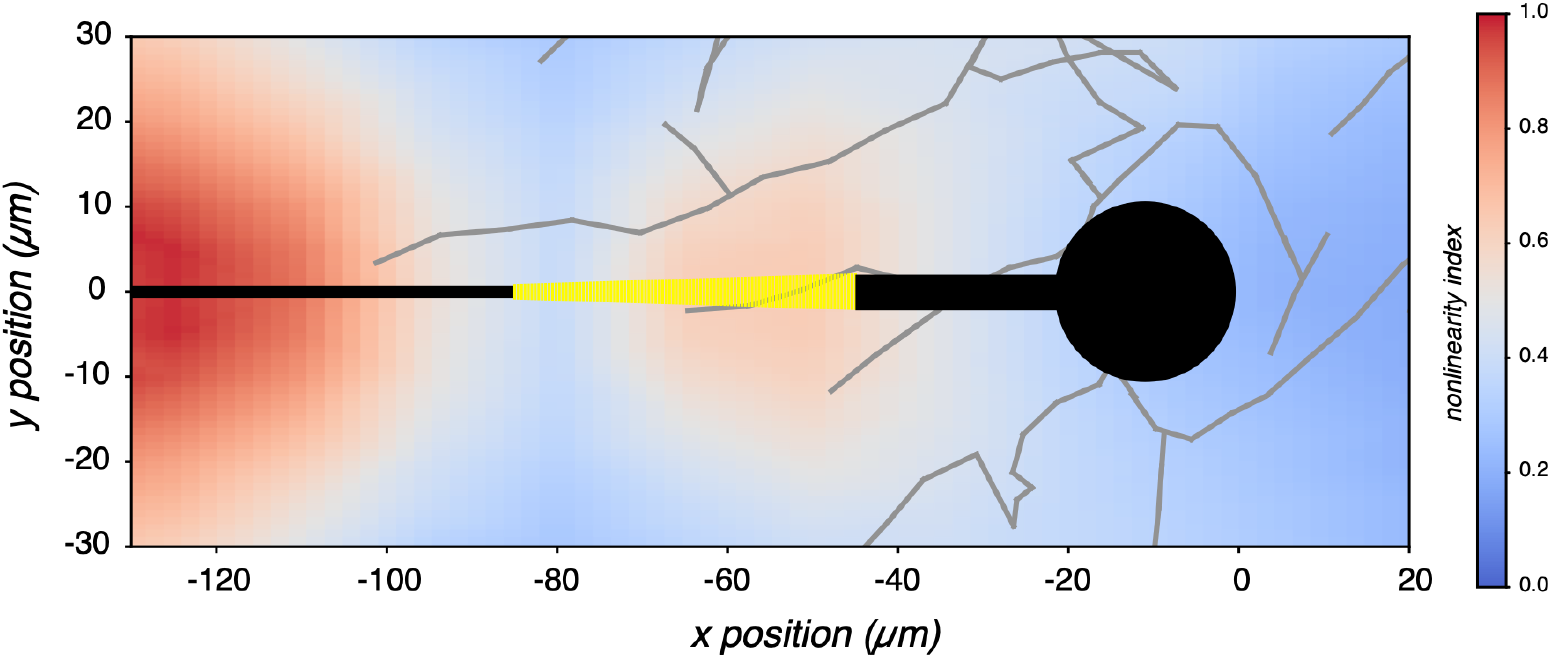
Response nonlinearity at somatic and dendritic regions. Colored according to the nonlinearity index (see Methods) for various placements of a two electrode stimulus around somatic and dendritic cell compartments. A nonlinearity index value of 1 (red) implies nonlinear current summations, while an index value of 0 (blue) implies linear current summations. Soma and axon shown in black, with the yellow portion highlighting the sodium channel band region of the axon. Dendrites shown in gray.

**Figure 6.**
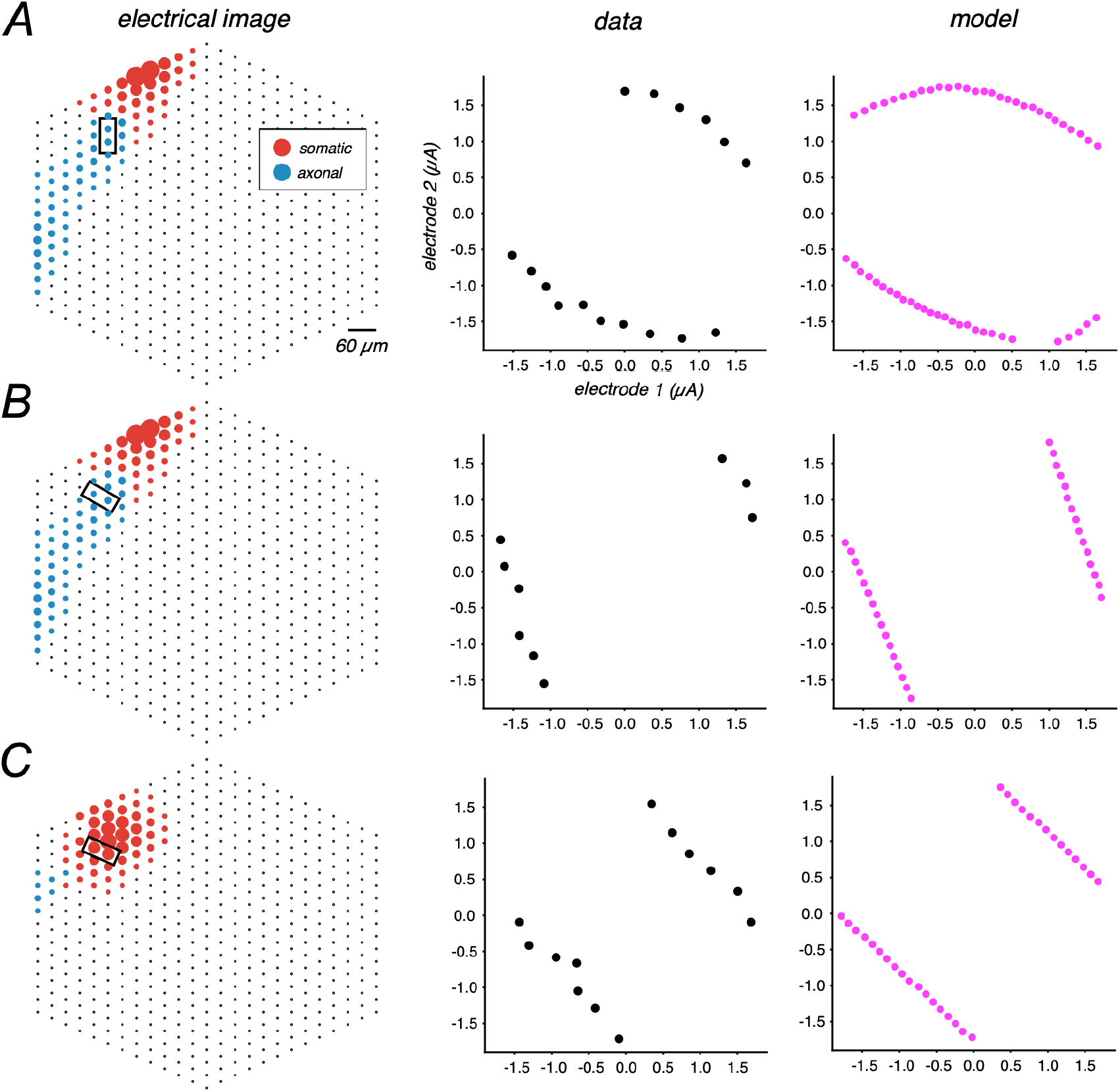
Comparing modeled and experimental responses to bi-electrode stimulation. ***A***, Left. Electrical image from an experimentally recorded parasol cell, circle area is proportional to recorded signal strength and color represents dominant compartment being recorded from (red: somatic; blue: axonal). Electrode pair oriented parallel to and positioned near the axon shown with a black rectangle. Middle. Experimentally recorded response, black points represent current combinations at threshold (0.5 probability). Right. Stimulated response profile for an electrode pair arrangement approximately matching the experimental condition, magenta points represent amplitude causing spiking for many fixed ratio current combinations that were tested. ***B*** and ***C***, similar to ***A***, but for electrode pairs positioned perpendicular to the distal axon and near the soma, respectively.

### Extension to three electrode stimulation

Similar trends were observed for simultaneous stimulation using three neighboring electrodes in a triangular arrangement, matching an arrangement used experimentally. Following a similar procedure as that for two electrode stimulation outlined above, the geometric arrangement of the collection of activation thresholds in the three-dimensional space of current levels was examined to test the linearity of summation. If currents combine linearly to drive response, this region should form a plane (or, for pulses with both positive and negative phases such as the ones used here, two planes on opposite sides of the origin).

For three-electrode stimuli placed close to the distal axon (**Fig. 7A, left**), the currents passed through the three electrodes generally combined nonlinearly to drive activation: the collection of thresholds formed a curved and closed shape in the space of electrical stimuli (**Fig. 7B, left**). This is consistent with the fact that regardless of the orientation of the three electrodes, multiple activation sites would be expected. In contrast, three-electrode placements far from the distal axon (>40 µm) or over the somatic and dendritic regions of the cell yielded more linear current summation (**Fig. 7A, middle & right**): the collection of thresholds approximately formed two planes in the space of electrical stimuli (**Fig. 7B, middle & right**). In the cases when summation was linear, the observed region of activation across different current levels was tightly localized (**Fig. 7C, middle & right**), while in the case of nonlinear summation, the region of activation was more spread out (**Fig. 7C, left**). Examination of current combinations at threshold revealed that in the nonlinear case the primary site of activation systematically shifted as the dominant current amplitude moved from one electrode to another (**Fig. 7B, left, colors**), whereas the dominant site was more stable in the linear cases **(Fig. 7B, middle & right, colors)**.

To experimentally verify these trends, the spiking behavior of parasol RGCs in response to three-electrode stimulation at various geometric placements was measured in *ex vivo* preparations of the macaque retina. The experimental trends broadly matched those seen in simulations. Specifically, three-electrode stimuli positioned close to the distal axon demonstrated nonlinear current summation (**Fig. 8A**), while stimuli positioned far (>40 µm) from the distal axon or over the somatic and dendritic regions of the cell yielded approximately linear current summation (**Fig. 8B-C**).

**Figure 7.**
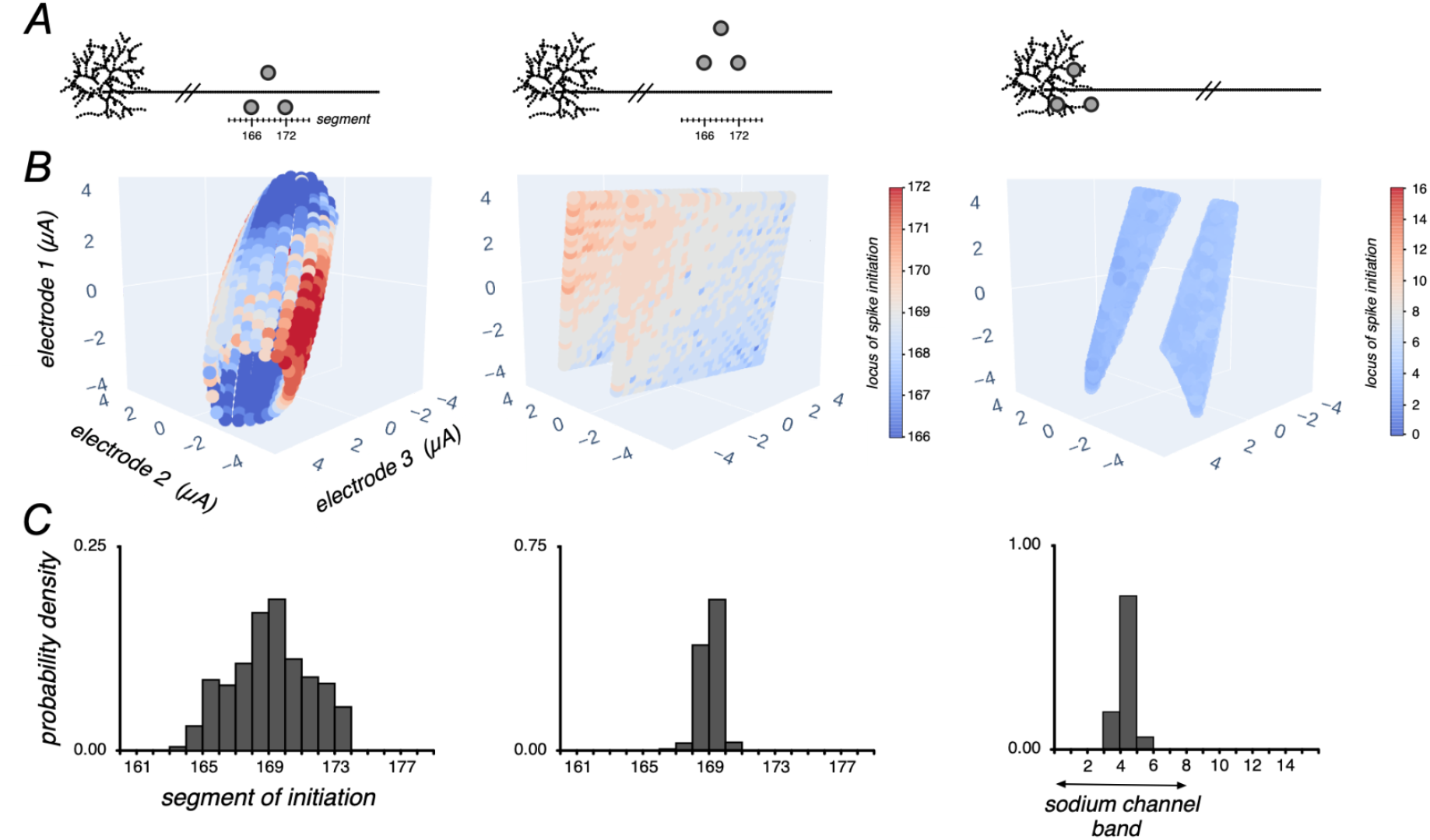
Responses to three electrode stimulation. ***A***, Visualization of the three electrodes in a triangular arrangement positioned at various locations along the model cell. The model cell is discretized into 5 µm length segments (see Methods). ***B***, Collection of activation thresholds in the three-dimensional space of current levels, colored by segment of spike initiation. ***C***, Distribution of spike initiation locations for the collection of current combinations causing activation that were colored in the middle panel (see Methods).

**Figure 8.**
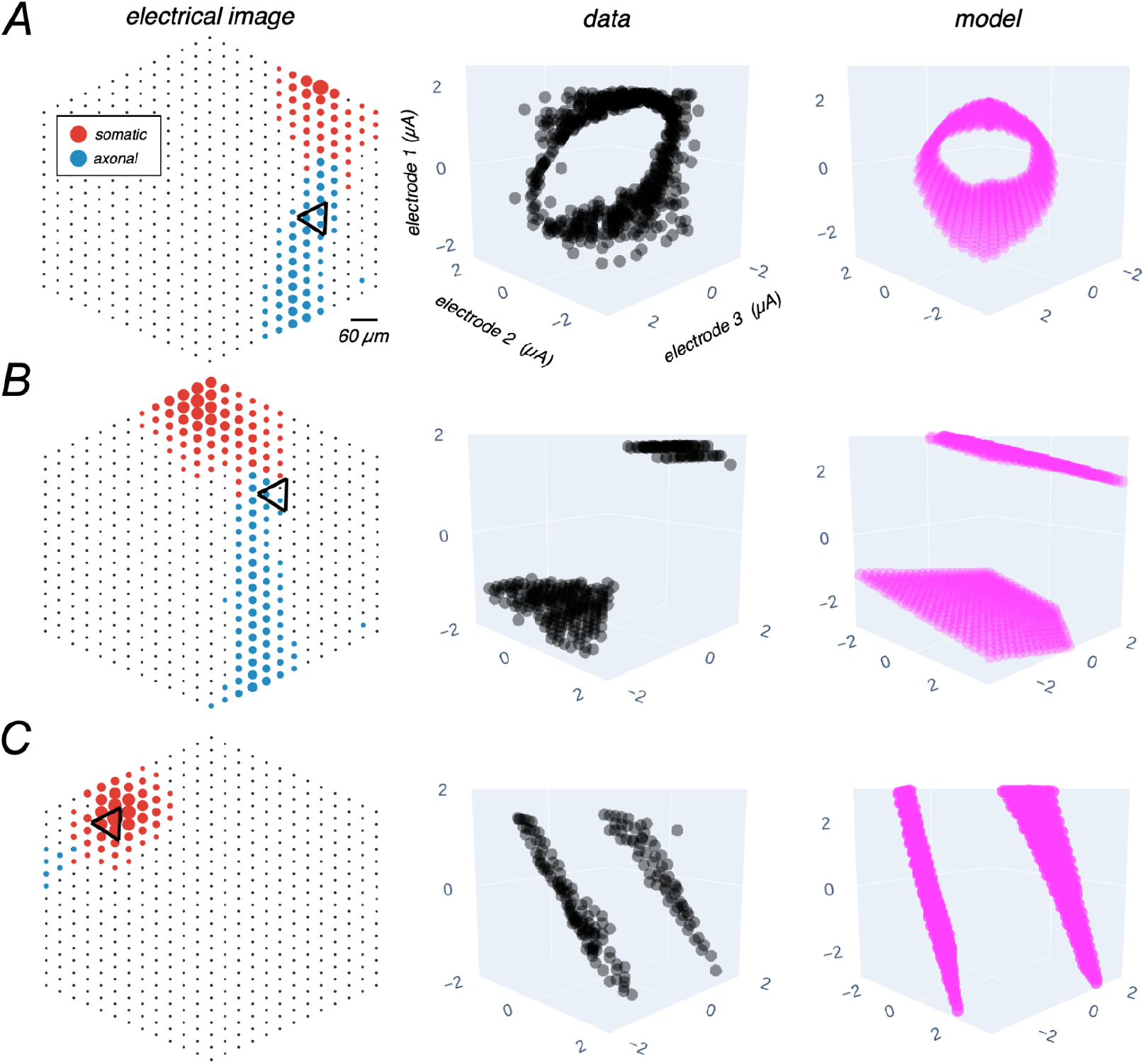
Comparing modeled and experimental responses to three-electrode stimulation. ***A***, Left. Electrical image from an experimentally recorded parasol cell, circle area is proportional to recorded signal strength and color represents dominant compartment being recorded from (red: somatic; blue: axonal). Three-electrode stimulus positioned near the axon, shown with a black triangle. Middle. Experimentally recorded response profile, black points represent current combinations yielding 0<p<1 probability spiking. Right. Stimulated response profile for a three-electrode arrangement approximately matching the experimental condition, magenta points represent amplitude causing spiking for many fixed ratio current combinations that were tested. ***B*** and ***C***, similar to ***A***, but for far axonal and somatic three-electrode stimulus placements, respectively.

## Discussion

We have presented a biophysical model (described schematically in **Fig. 1)** capable of reproducing individual RGC responses to multi-electrode stimulation, validated it against experimental data, and tested the multi-site stimulation hypothesis for response nonlinearity. The model reproduced essential features of single-electrode recording and stimulation (**Fig. 2**) and the experimentally observed linearity and nonlinearity of current summation for stimulation using multiple simultaneously active electrodes (**Fig. 6**,**8**). The positioning of the multi-electrode stimulus relative to the cell was identified as a key factor affecting the response nonlinearity (**Fig. 3, 4, 5, 7; A**), including the orientation of electrodes relative to the axon (**Fig. 3**) and the distance from the axonal (**Fig. 4**) and somatic/dendritic regions (**Fig. 5**) of the cell. The response nonlinearity increased with the number of distinct initiation sites accessed by the multi-electrode stimuli (**Fig. 3, 4, 7; B,C**). All of these features of the simulations closely paralleled experimental observations **(Fig. 6, 8)**. These findings support multi-site activation as the biophysical mechanism driving nonlinear RGC responses and highlight the value of modeling to understand how implanted devices can be used to modulate neural activity.

The multi-site activation hypothesis accounts for nonlinear responses to multi-electrode stimulation as follows. Spike initiation at one site on a cell is governed by the transmembrane voltage, which is determined by a linear combination of currents applied through the multiple electrodes. However, as the ratio of currents passed through the electrodes is changed, the site of spike initiation shifts. At the new site, the linear relationship between current and activation is different because its position relative to the two electrodes is different. Thus, the shift to a new site of spike initiation produces an overall nonlinear response as a function of multi-electrode currents. The present simulation results and empirical tests are consistent with this biophysical explanation of nonlinear activation.

The close agreement between model predictions and experimental data highlights the usefulness of the biophysical model. Ion channel dynamics from previous work [14] were used to define the electrical properties of our RGC cable model. These dynamics have been employed in many other epiretinal stimulation modeling studies [15,17,30,31]. The present findings show that the dynamics accurately capture complex RGC response behaviors with various cell-electrode geometries, which demonstrates its capability and supports its use as a tool to better understand RGC firing and responses to electrical stimulation. Note, however, that action potential dynamics in the macaque retina were not fully explained by the model (see below).

No attempt was made to quantitatively match the stimulation thresholds and recorded spike waveforms between the data and model on every electrode for every cell. Instead, we showed that a standard multi-compartment model produced important trends in both recorded spikes and evoked spikes that were qualitatively consistent with those observed experimentally. This consistency supported the use of the model to probe mechanisms underlying multi-electrode stimulation and better understand factors affecting linear/nonlinear RGC response. However, future efforts to quantitatively match experimental data in detail could yield additional insights.

Previous studies have demonstrated examples in which current passed through multiple electrodes combines nonlinearly to drive neural response, with the most direct evidence coming from studies targeting single RGCs in the retina [4,6]. This nonlinearity makes the evoked responses difficult to predict and control, and therefore limits the practical use of multi-electrode stimulation strategies to improve targeting of cells. This difficulty was accounted for in the previous work by fitting linear-nonlinear or piecewise linear models, leveraging the multi-site activation hypothesis as a justification [4,6]. However, unlike the present work, these studies did not provide a test of the multi-site hypothesis nor did they provide a way to predict the degree of nonlinearity based on the spatial arrangement of the cell and electrodes.

Some studies have also suggested that multi-site activation is predicted by the activating function (AF), which is proportional to the second spatial derivative of extracellular voltage along the membrane for a homogeneous fiber with constant diameter [7,32]. In this approximation, different current combinations through multiple electrodes result in distinct peaks in the AF, implying potentially distinct loci of activation. However, the AF is an inaccurate predictor of thresholds and spike initiation locations for non-infinite model axons [8,9]. Specifically, while the AF provides a prediction of the region of membrane undergoing the greatest depolarization in response to stimulation, the prediction is only accurate instantaneously after stimulus onset.

Dispersion of charge immediately after stimulus onset causes a wider region of depolarization, diminishing the predictive power of the AF. Additionally, finite axon geometries and inhomogeneity in the ion channel density along the axon result in non-uniform current flows within the membrane, further limiting the usefulness of the AF, which assumes an infinitely long axon with uniform current flow. Thus, although the AF is conceptually useful, the biophysical simulation provides a more direct and accurate explanation of multi-site activation.

Several caveats of the present work deserve mention.

(1) The model was developed to mimic experimental parameters from a laboratory prototype epiretinal implant device (512 electrodes, 30 µm pitch, 5-10 µm diameter) [12]. This allowed for experimental validation of the model. Other epiretinal devices often use larger diameter electrodes (100-200 µm), with larger spacing between electrodes (>100 µm), positioned farther from the surface of the retina (>100 µm) [1,3]. Evidence of linear summation has been observed with such a device (Argus II) when using paired electrode stimulation abiding by certain placement geometries relative to the axons, indicating that similar principles may apply [33]. However, these MEA geometries would need to be tested in the model to fully understand the biophysical causes of linear and nonlinear summation in electrically evoked responses using those devices.
(2) Simplifying assumptions were made when modeling the electrical properties of the extracellular environment. Specifically, the extracellular medium was defined to have an isotropic resistivity of 1000 Ohm-cm. In contrast, the real extracellular medium is a non-isotropic medium, with previous work describing different resistivity values in different retinal layers [34]. However, for electrode arrays positioned close to the RGC layer (<10 µm), isotropic resistivity is likely a reasonable assumption. Additionally, the extracellular voltage profiles elicited by the disk electrodes were approximated using analytical expressions [16]. While both of these simplifications enable computational efficiency, future work using more precise finite-element modeling of both the extracellular medium and electrodes may be useful.
(3) This work exclusively explored biophysical simulation of RGCs, specifically ON and OFF parasol cells, which are the most reliably recorded and stimulated cell types in the experimental preparations used to validate the model. Two other numerically dominant cell types in the primate retina are the ON and OFF midget cells, which convey high spatial resolution information to the brain. Additional work must be done to understand how the presented findings apply to these cell types, which differ from parasol cells in their morphology and biophysical properties [35]. Beyond the retina, neural stimulation devices might be used to interface with neurons with properties very different from RGCs, such as myelinated axons. To explore the generalizability of the present findings for other brain regions, it would be necessary to model those specific neuron types.
(4) The temporal dynamics of the modeled spikes were slower than what was observed experimentally (**Fig. 2, top**). First, the spike width of the depolarization phase of the modeled action potential is larger than that of experimentally observed action potentials. This discrepancy could be due to a difference in the animal model: the data shown were obtained from primate RGCs, whereas the biophysical parameters were obtained from a study done with rat and cat RGCs [14]. Second, the hyperpolarizing phase of the modeled action potential is much slower than what is observed experimentally. Recent modeling work has incorporated hyperpolarizing activated currents (*I* _*h*_), which could increase the speed at which the modeled spike returns to the resting potential [35].
(5) The present analysis focused on investigating responses to short charge-balanced stimulation pulses, specifically, triphasic pulses matching those used experimentally. Similar trends held for more commonly used symmetric biphasic stimulation pulses (data not shown). While less common, long asymmetric charge-balanced stimulation pulses have also been used [36]. Additional analysis would be necessary to explore the generalizability of the present findings for such stimuli.

The present results provide geometric guidelines for predicting the linearity of current summation, which could be used for the design of future multi-electrode stimulation strategies. Electrodes positioned perpendicular to the axon or at any orientation over the cell body demonstrate approximately linear current summation, suggesting the following experimental approach. First, features from electrical recording (e.g., EIs) could be used to determine the axon trajectories of all recorded RGCs, which tend to be parallel in a given region of the retina. Second, activation thresholds from single-electrode stimulation could be used to determine the influence of individual electrodes on the activation of nearby RGCs. Then, a collection of stimulating electrodes arranged perpendicular to the axons would be chosen to ensure linear summation of current passed simultaneously through any subset of these electrodes. The predictable linearity of such a stimulus combined with the known single-electrode thresholds could then be exploited by algorithms to efficiently design multi-electrode stimulation patterns for more targeted RGC activation. Therefore, the observed geometric guidelines for linearity provide a strong prior that could enable the more effective design of multi-electrode strategies for use in future devices.

## Acknowledgments

This work was supported by the Stanford Graduate Fellowship and the Stanford Bio-X Bowes Fellowship (RSV), Research to Prevent Blindness Stein Innovation Award, Wu Tsai Neurosciences Institute Big Ideas, NIH NEI R01-EY021271, and NIH NEI P30-EY019005 (EJC), Polish National Science Centre Grant DEC-2013/10/M/NZ4/00268 (PH), AGH UST, task No. 11.11.220.01/4 within subsidy of the Ministry of Science and Higher Education (WD). We thank Paul Werginz, Jim Weiland, Ted Carnevale, and Sasi Madugula for useful discussions.

